# Global prevalence of naturally occurring *Wolbachia* in field-collected *Aedes* mosquitoes: a systematic review and meta-analysis

**DOI:** 10.1101/2024.09.19.614020

**Authors:** Tewelde T. Gebremariam, Polly Leung, Vincent Rusanganwa

## Abstract

**Background:** Dengue is one of the leading causes of morbidity worldwide. *Wolbachia-*mediated dengue biocontrol has emerged as a novel strategy in recent decades and depends on a lack of *Wolbachia* in the natural populations of *Aedes* mosquitoes. Through a systematic review of the published literature, this study sought to estimate the natural occurrence of *Wolbachia* among field-collected *Aedes* mosquitoes worldwide.

**Methods:** We conducted hand and systematic searches from PubMed, the Cochrane Library, and Google Scholar for all relevant published articles on *Wolbachia* infection in *Aedes* mosquitoes published before September 12, 2024. The prevalence estimates were analysed using a random effects meta-analysis, and a random effects meta-regression was performed to determine sources of heterogeneity in prevalence estimates.

**Results:** Twenty-three studies with 16,997 field-collected *Aedes* mosquitoes from different countries worldwide were included. The meta-analysis revealed a global pooled prevalence of natural *Wolbachia* infection in *Aedes* mosquitoes of 57.7% (95% CI: 41.0–72.8%), which was six times higher in *Ae. albopictus* than *Ae. aegypti (p* <0.001) and ranged from 6.0% (95% CI: 2.6–13.1%) in *Ae. aegypti* to 87.1% (95% CI: 78.0–92.8%) in *Ae. albopictus*. Continentally, Asia had the highest percentage of *Wolbachia* infection in *Ae. aegypti* (7.1%), followed by Europe (5.0%), North America (1.9%), and Africa (0.7%). Similarly, Asia had the highest prevalence of *Wolbachia* in *Ae. albopictus* (95.5%), followed by Europe (94.8%), North America (91.6%), South America (85.2%), and Africa (71.6%). Overall, dengue incidence was negatively related to *Wolbachia* prevalence (B = −0.0006, *p* = 0.0281). Species wise, infection rates in *Ae. aegypti* were significantly higher among females than males (OR = 1.72; 95% CI = 1.01, 2.92, *p* = 0.046), while there was no difference between males and females in *Ae. albopictus* (*p* = 0.098). Furthermore, *Wolbachia* infection rates in *Ae. albopictus* were inversely correlated with dengue incidence (β = −0.0013, p <0.01) but not in *Ae. aegypti (p =* 0.0984). In contrast, higher temperature was negatively associated with *Wolbachia* prevalence in *Ae. aegypti* but not in *Ae. albopictus*. In contrast, higher temperature was negatively associated with *Wolbachia* prevalence in *Ae. aegypti* (β = --2.5736, p <0.001) but not in *Ae. albopictus (p =* 0.7050).

**Conclusion:** *Aedes* mosquitoes had a high and variable prevalence of naturally occurring *Wolbachia*, and this was negatively correlated with dengue incidence across countries. While the natural infection of *Ae. albopictus* is more common, detection in *Ae. aegypti* may reflect contamination and require high-quality multicentre studies to verify the above findings.

## Background

Dengue is a major public health concern, and the incidence has dramatically increased since approximately 1990 (1). There are currently an estimated 100–400 million dengue cases, with around half the world’s population at risk of acquiring dengue (2). The disease is transmitted through the bite of female *Aedes* mosquitoes, specifically *Aedes aegypti* and *Ae. albopictus*, infected by either of the four antigenically distinct dengue virus serotypes (DENV-1, -2, -3, and -4) (3). Traditional *Aedes* mosquito control strategies, such as chemical sprays are unable to stop the spread of DENVs and have instead caused insecticide resistance to emerge, necessitating the development of novel approaches (4, 5).

Because of their low fitness costs, ability to generate cytoplasmic incompatibility (CI), high rates of maternal transfer, and viral blocking effects, *Wolbachia* have gained popularity in recent decades in reducing dengue transmission (6-8). *Wolbachia* species are widespread noncultivable, obligate intracellular gram-negative endosymbiotic bacteria that belong to the *Alphaproteobacterial* class, Rickettsiales order, and genus *Wolbachia* and infect most arthropods and some filarial nematodes (9, 10). *Wolbachia* between hosts is primarily maternally inherited, although horizontal transmission can also occur. Vertical transmission occurs when *Wolbachia*-carrying female insects pass through the endosymbiont via their eggs to their offspring, thus generating a stable line of *Wolbachia*-insect interaction. In horizontal transmission, the bacteria rapidly spread between insect populations and colonise new species (11). Approximately 25-70% of insect species are naturally infected with *Wolbachia*, making it one of the most abundant endosymbionts on Earth (12, 13).

Classified as a single species, *Wolbachia pipientis* is grouped into 16 supergroups or clades, denoted A–F and H–Q, with supergroups A and B exclusively infecting terrestrial arthropods, including vectors transmitting malaria and arboviruses (10, 14). While *Ae. albopictus* is a natural *Wolbachia* host (15), reports of *Ae. aegypti* infections have surfaced recently (16-31). The tissue distribution of *Wolbachia* infections varies greatly; native strains are usually limited to the germline tissues, whereas nonnative strains typically exhibit a wider range of tissues and higher concentrations in somatic tissues, which is particularly relevant for vector control (32-34).

Through a process known as CI, *Wolbachia* alters the host cell environment, affecting host reproduction. Males infected with Wolbachia become sterile when they mate with uninfected females but not infected females (rescue mating) (35-37). To survive intracellularly, *Wolbachia* uses the type IV secretion system and its secreted effector proteins that modulate bacteria-host interactions and sculpt the cellular environment to persist, replicate, and be maternally transmitted (38). The mechanisms of DENV inhibition appear to be multifactorial and include the induction of endoplasmic reticulum stress and reactive oxygen species (39, 40); activation of the host cell innate immunity (41), alteration of the host actin cytoskeleton (42); perturbation of endocytosis, vesicular trafficking, lipid metabolism, and; disruption of endoplasmic reticulum (43-45); and dysregulation of RNA-binding proteins, such as XRN1 and DNMT2 (46-48). Viral escape mutations that restore viral replication are less likely to arise because multiple cellular disruptions are more likely to contribute to viral inhibition (32).

The ability of *Wolbachia* to impede virus replication was initially demonstrated in *Drosophila melanogaster*, and the *Wolbachia* that passed through the maternal line were able to shield these insects against the Drosophila C virus (49, 50), a positive-stranded RNA virus that kills fruit flies within 3–4 days (51). The genome of the virus was also important since *Wolbachia* was found to be protective against RNA viruses but not DNA viruses (49). Viruses blocked by *Wolbachia* tend to have positive-stranded RNA genomes, which include a vast majority of arboviruses, including DENV (32).

The concept of spreading virus-blocking *Wolbachia* to reduce dengue transmission, which includes population suppression and population replacement, has gained increased attention in recent years. The population replacement strategy involves releasing females infected with *Wolbachia* after mating with males (*Wolbachia*-infected or not), producing viable offspring, and achieving a high frequency of *Wolbachia* establishment in the population, thereby reducing the transmission competency of that population while keeping the overall number of mosquitoes unchanged. The population suppression strategy, however, involves the release of male *Wolbachia*-infected mosquitoes, which reduces the total population of mosquitoes by preventing them from producing viable offspring when they mate with wild females (52, 53).

The strains of *Wolbachia*, the interaction of different *Wolbachia* strains and other bacterial community members, and the genetic makeup of *Ae. aegypti* influence how successful *Wolbachia*-mediated dengue control is (54). *Wolbachia* strains vary considerably in their effects on *Ae. aegypti* vector competency and viral blocking, fitness costs, and temperature stability (55). The ideal traits for *Wolbachia* strains under such strategies include high levels of viral inhibition, strong penetrance of the CI phenotype, high rates of maternal transmission, minimal fitness costs, and environmental stability (32).

Generation of *Wolbachia*-infected *Ae. aegypti* lines has been done with several different *Wolbachia* strains, including *wAlbA, wAlbB, wAu*, and w*Mel. wAlbA* and *wAlbB* are both native to *Ae. albopictus*, while *wAu* and w*Mel* are derived from *Drosophila simulans* and *D. melanogaster*, respectively (56, 57). wAlbB showed very strong CI induction, good viral blocking, and high fitness. This appears to vary with the host genotype. Critically, however, wAlbB proved to be much less susceptible to the effects of high rearing temperatures than wMel, and wAlbB may be well suited for population replacement in a very hot environment (57).

*Wolbachia* infection has been successful at reducing the incidence of dengue in some areas. A trial has shown that infection of *Ae. aegypti* with wMel-*Wolbachia* decreases the probability of contracting virologically confirmed dengue by 77% (6). Nevertheless, the antagonistic interactions between naturally occurring and experimentally infected *Wolbachia* in the target population could jeopardise the efficacy of both population replacement and suppression programmes (58). Furthermore, the fitness deficiencies observed in the artificial system currently employed in *Wolbachia* release programmes, which most likely restrict the spread of *Wolbachia* after mosquito release, could be overcome by the naturally existing *Wolbachia-Aedes* association (59). These approaches may also be more successful at mitigating arboviral diseases if they are applied in areas with no evidence of natural *Wolbachia* infections in *Ae. aegypti* mosquitoes (60). The variable occurrence of *Wolbachia* strains in *Ae. aegypti* across different habitats could eventually have an impact on arbovirus-focused biocontrol initiatives. Consequently, determining the natural occurrence, genetic diversity, and geographic distribution of *Wolbachia* is central for both planning and implementing *Wolbachia* release operations (61). The purpose of this review was to gather evidence on the pooled prevalence of natural *Wolbachia* infection in *Aedes* mosquitoes and how this is related to dengue incidence.

## Methods

The Preferred Reporting Items for Systematic Reviews and Meta-Analyses (PRISMA) checklist (62) was used as part of the review methodology. The methodological quality of the included studies was evaluated using the Joanna Briggs Institute (JBI) Critical Appraisal Checklist for Studies Reporting Prevalence Data (63).

### Inclusion and exclusion criteria

Cross-sectional studies that reported natural *Wolbachia* infections in *Ae. aegypti* and *Ae. albopictus* were included. *Wolbachia* might be detected by conventional polymerase chain reaction (PCR), nested PCR (nPCR), quantitative PCR (qPCR), or loop-mediated isothermal amplification (LAMP) assays.

Studies with a small sample size (n <30); reviews; grey literature; books; posters; conference proceedings; unpublished articles; studies on artificial infection; publications whose full texts were not available in English, articles based on previously published samples; experiments; and studies reporting the presence of *Wolbachia* in species other than *Ae. aegypti* were excluded. The primary outcome measure was the prevalence of natural *Wolbachia* in *Aedes* mosquitoes.

### Databases and search strategy

The review question was formulated using the CoCoPop mnemonic, which stands for “condition, context, and population” (63). The PubMed, Google Scholar, and Cochrane databases were systematically searched from inception to September 12, 2024. The following was the search approach employed:

### PubMed

((“*Wolbachia*”[Mesh] OR *Wolbachia*[tiab]) AND (“*Aedes*”[Mesh] OR *Aedes*[tiab])) OR ((*Wolbachia*[tiab] AND *Aedes*[tiab]) NOT MEDLINE[sb]) NOT systematic[sb]

### Cochrane library

Cochrane library:
| ID | Search |
| --- | --- |
| #1 | MeSH descriptor: [Wolbachia] explode all trees |
| #2 | ("Wolbachia"):ti,ab,kw (Word variations have been searched) |
| #3 | MeSH descriptor: [*Aedes*] explode all trees |
| #4 | ("Aedes aegypti"):ti,ab,kw OR ("Aedes albopictus"):ti,ab,kw (Word variations have been searched) |
| #5 | #1 OR #2 |
| #6 | #3 OR #4 |
| #7 | #5 AND #6 |

### Study quality appraisal and data extraction

Two authors (TG and VR) independently screened each article at the abstract and full-text levels. Discrepancies between the two reviewers were resolved through discussion. Articles endorsed in the full-text screening were subjected to the 9-point Joanna Briggs Institute critical appraisal tool (63). We extracted the data from the selected studies using Microsoft Excel. The following data were extracted: the last name of the first author, year of publication, country of origin, dengue incidence, sample size, mosquito host sex, number of *Wolbachia* infections, supergroups, detection methods, and molecular markers. Studies were deemed to be of sufficient quality for inclusion in the meta-analysis if their JBI score was four or higher.

### Statistical analysis

The prevalence estimates were examined using a random effects meta-analysis. The I^2^ statistic and the Cochran’s Q-test were used to evaluate the heterogeneity of the included studies. To determine the sources of heterogeneity, random effects meta-regression was employed. The Begg and Mazumdar rank correlation tests were used to examine publication bias. The prevalence of *Wolbachia* by *Aedes* mosquito species and continent was examined using a subgroup analysis, and the effect of host sex was estimated by computing the odds ratio (OR) and its 95% confidence interval (CI). A p-value of less than 0.05 was deemed to be statistically significant. The Spearman’s test examined correlations between dengue incidence and *Wolbachia* infection in *Aedes* mosquitoes. Comprehensive Meta-analysis (version 3) was used to analyse the data.

### Patient and public involvement

This study was performed without patient or public involvement.

## Results

### Literature retrieval and characteristics of the included studies

The PRISMA flowchart (Fig. **1**) shows the article screening and selection process. After 1,218 records were identified via the databases’ literature retrieval process, the titles and abstracts of 1,114 of those records were carefully examined. The complete texts of 36 studies (15, 17, 18, 20, 21, 24-26, 56, 59, 61, 64-88), including 16,997 field-collected *Aedes* mosquitoes (7,531 *Ae. aegypti* and 9,466 *Ae. albopictus*), were included in the quantitative synthesis (Table **1**). Of the studies included, nineteen were conducted in Asia (15, 20, 21, 24-26, 56, 64, 67-71, 73-75, 79, 81, 85), five in Africa (66, 72, 78, 84, 88), four in Europe (61, 80, 83, 87), three on more than one continent (17, 59, 86), three in North America (18, 76, 77), and two in South America (65, 82).

**Fig. 1.**
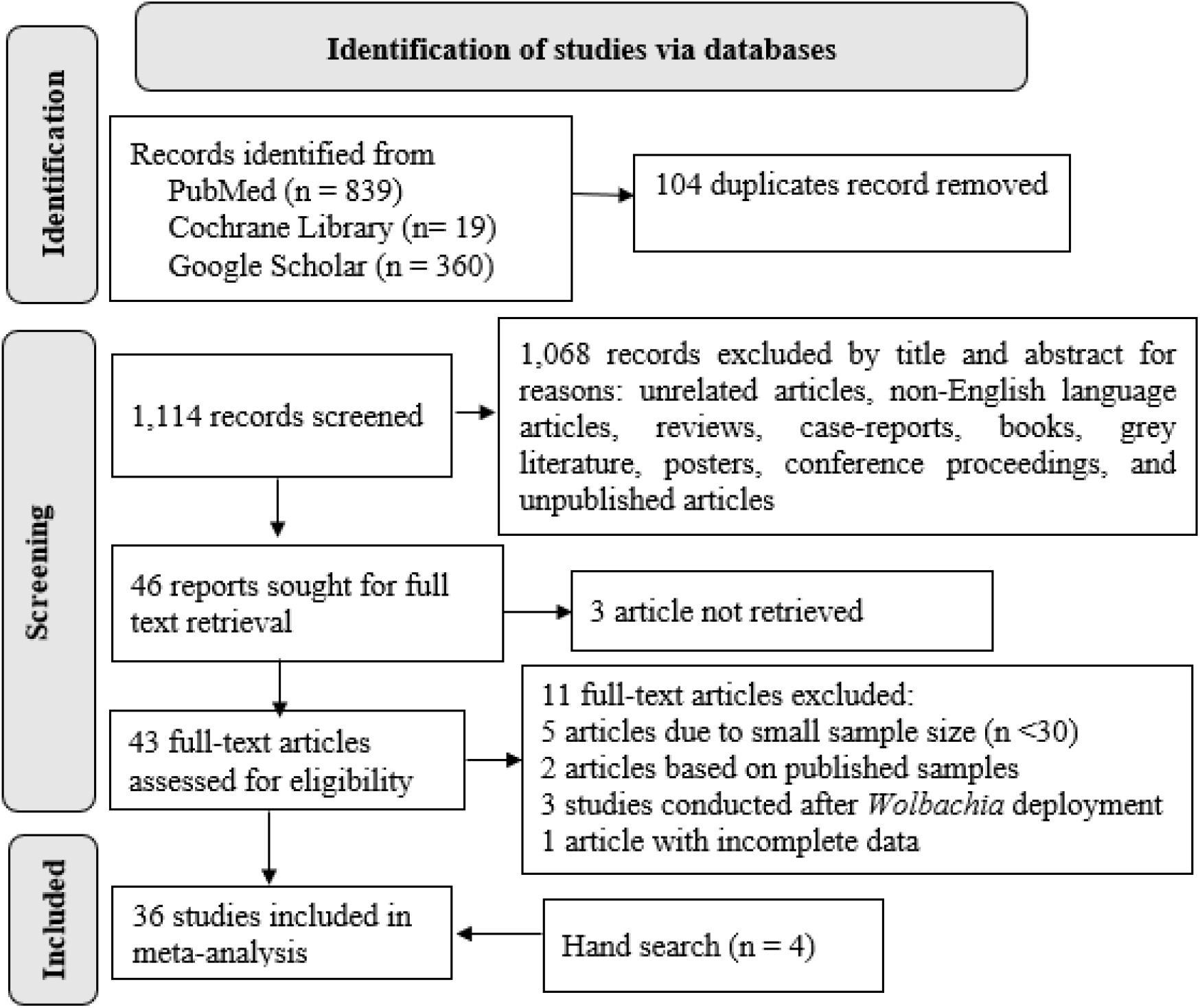
Flow diagram for study screening and selection process

The *Wolbachia* detection methods employed in the included studies were conventional PCR in 28 studies (15, 18, 21, 24, 26, 56, 59, 61, 64-69, 71-77, 79, 82-85, 88, 89); qPCR in five studies (29, 78, 80, 81, 86); multiplex PCR in two studies (70, 87); nPCR in two studies (20, 25); and LAMP in one study (17) (Table 1). The targeted *Wolbachia* genes were *wsp* in 21 studies (18, 21, 24-26, 56, 66-74, 77, 79, 80, 82, 84, 85); 16S rRNA in seven studies (17, 18, 20, 61, 76, 78, 86); 16S rDNA in two studies (15, 75); *wsp* and 16S rRNA in three studies (81, 83, 88), *gatB* or *ftsZ* in one study (17); and *SVNP* and *wsp* in one study (64) (Table **1**).

**Table 1.**
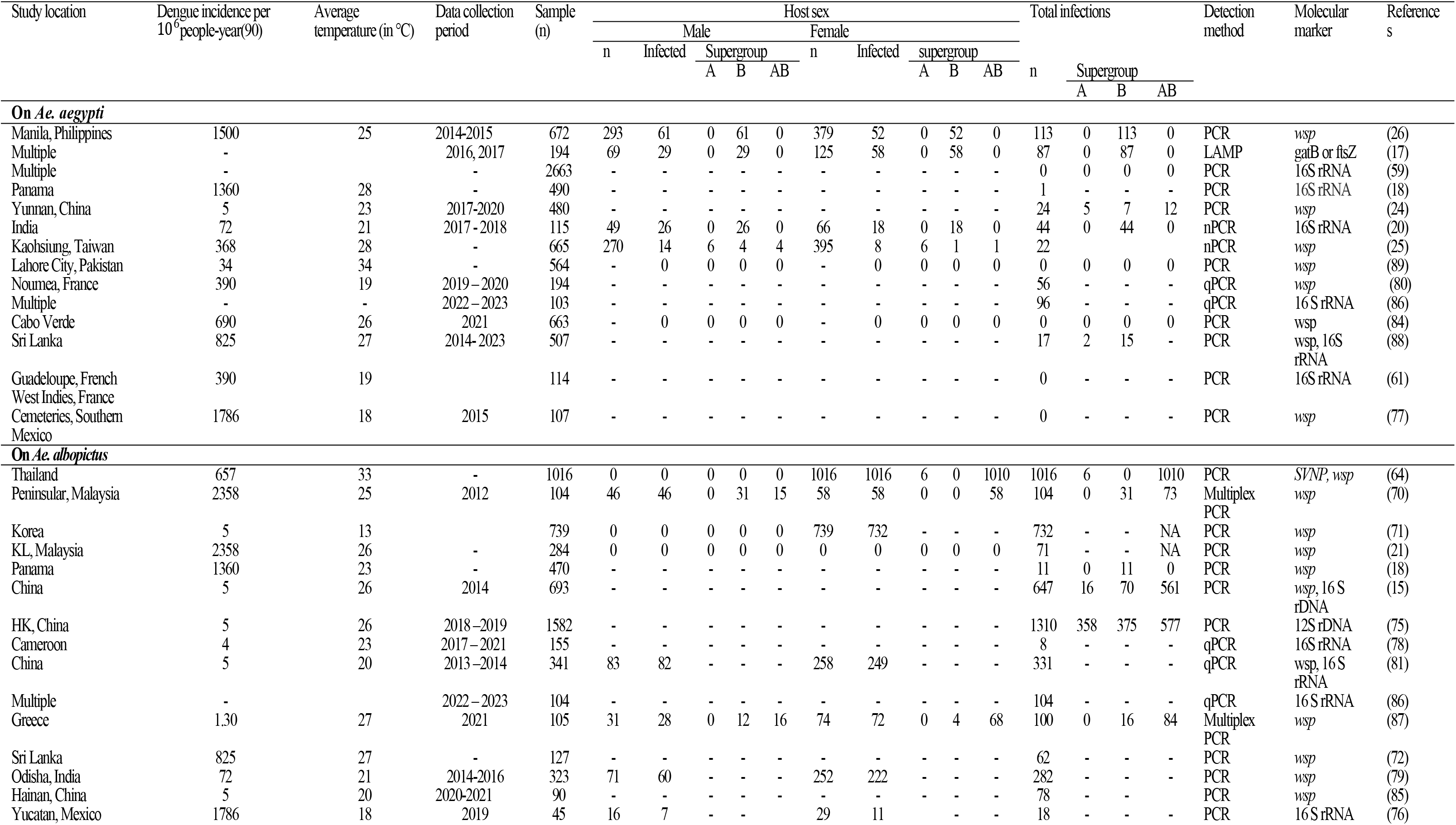
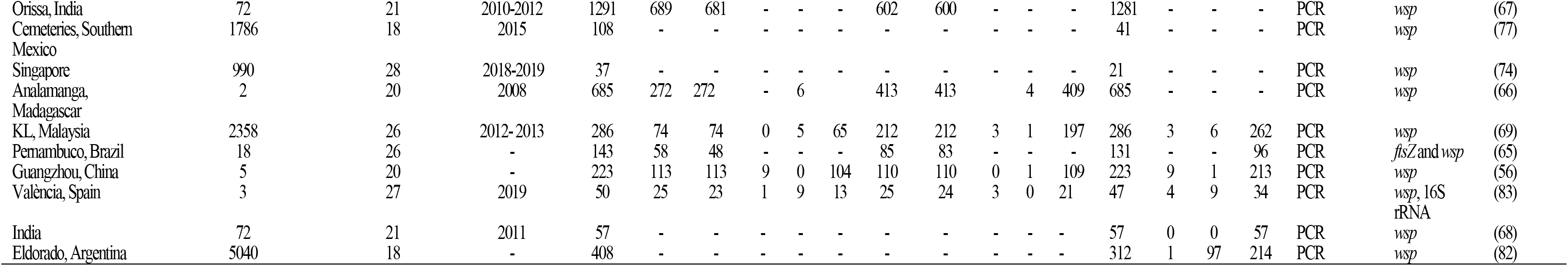
Characteristics of the included studies on *Aedes* mosquitoes.

| Study location | Dengue incidence per 10<sup>6</sup>people-year(90) | Average temperature (in °C) | Data collection period | Sample (n) | Host sex |  |  |  |  |  |  |  |  | Total infections |  |  | Detection method | Molecular marker | Reference s |  |  |
| --- | --- | --- | --- | --- | --- | --- | --- | --- | --- | --- | --- | --- | --- | --- | --- | --- | --- | --- | --- | --- | --- |
|  |  |  |  |  | Male |  |  |  |  | Female |  |  |  |  |  |  |  |  |  |  |  |
|  |  |  |  |  | n | Infected | Supergroup |  |  | n | Infected | supergroup |  |  |  |  |  |  |  |  |  |
|  |  |  |  |  |  |  | A | B | AB |  |  | A | B | AB | n | Supergroup |  |  |  |  |  |
|  |  |  |  |  |  |  |  |  |  |  |  |  |  |  |  |  | A | B | AB |  |  |
| ***On Ae. aegypti*** |  |  |  |  |  |  |  |  |  |  |  |  |  |  |  |  |  |  |  |  |  |
| Manila, Philippines | 1500 | 25 | 2014-2015 | 672 | 293 | 61 | 0 | 61 | 0 | 379 | 52 | 0 | 52 | 0 | 113 | 0 | 113 | 0 | PCR | *wsp* | (26) |
| Multiple | - |  | 2016, 2017 | 194 | 69 | 29 | 0 | 29 | 0 | 125 | 58 | 0 | 58 | 0 | 87 | 0 | 87 | 0 | LAMP | *gatB* or *ftsZ* | (17) |
| Multiple | - |  | - | 2663 | - | - | - | - | - | - | - | - | - | - | 0 | 0 | 0 | 0 | PCR | 16S rRNA | (59) |
| Panama | 1360 | 28 | - | 490 | - | - | - | - | - | - | - | - | - | - | 1 | - | - | - | PCR | 16S rRNA | (18) |
| Yunnan, China | 5 | 23 | 2017-2020 | 480 | - | - | - | - | - | - | - | - | - | - | 24 | 5 | 7 | 12 | PCR | *wsp* | (24) |
| India | 72 | 21 | 2017 - 2018 | 115 | 49 | 26 | 0 | 26 | 0 | 66 | 18 | 0 | 18 | 0 | 44 | 0 | 44 | 0 | nPCR | 16S rRNA | (20) |
| Kaohsiung, Taiwan | 368 | 28 | - | 665 | 270 | 14 | 6 | 4 | 4 | 395 | 8 | 6 | 1 | 1 | 22 |  |  |  | nPCR | *wsp* | (25) |
| Lahore City, Pakistan | 34 | 34 | - | 564 | - | 0 | 0 | 0 | 0 | - | 0 | 0 | 0 | 0 | 0 | 0 | 0 | 0 | PCR | *wsp* | (89) |
| Noumea, France | 390 | 19 | 2019–2020 | 194 | - | - | - | - | - | - | - | - | - | - | 56 | - | - | - | qPCR | *wsp* | (80) |
| Multiple | - | - | 2022–2023 | 103 | - | - | - | - | - | - | - | - | - | - | 96 | - | - | - | qPCR | 16 S rRNA | (86) |
| Cabo Verde | 690 | 26 | 2021 | 663 | - | 0 | 0 | 0 | 0 | - | 0 | 0 | 0 | 0 | 0 | 0 | 0 | 0 | PCR | *wsp* | (84) |
| Sri Lanka | 825 | 27 | 2014- 2023 | 507 | - | - | - | - | - | - | - | - | - | - | 17 | 2 | 15 | - | PCR | *wsp*, 16S rRNA | (88) |
| Guadeloupe, French West Indies, France | 390 | 19 |  | 114 | - | - | - | - | - | - | - | - | - | - | 0 | - | - | - | PCR | 16S rRNA | (61) |
| Cemeteries, Southern Mexico | 1786 | 18 | 2015 | 107 | - | - | - | - | - | - | - | - | - | - | 0 | - | - | - | PCR | *wsp* | (77) |
| ***On Ae. albopictus*** |  |  |  |  |  |  |  |  |  |  |  |  |  |  |  |  |  |  |  |  |  |
| Thailand | 657 | 33 | - | 1016 | 0 | 0 | 0 | 0 | 0 | 1016 | 1016 | 6 | 0 | 1010 | 1016 | 6 | 0 | 1010 | PCR | *SNP*, *wsp* | (64) |
| Peninsular, Malaysia | 2358 | 25 | 2012 | 104 | 46 | 46 | 0 | 31 | 15 | 58 | 58 | 0 | 0 | 58 | 104 | 0 | 31 | 73 | Multiplex PCR | *wsp* | (70) |
| Korea | 5 | 13 |  | 739 | 0 | 0 | 0 | 0 | 0 | 739 | 732 | - | - | - | 732 | - | - | NA | PCR | *wsp* | (71) |
| KL, Malaysia | 2358 | 26 | - | 284 | 0 | 0 | 0 | 0 | 0 | 0 | 0 | 0 | 0 | 0 | 71 | - | - | NA | PCR | *wsp* | (21) |
| Panama | 1360 | 23 | - | 470 | - | - | - | - | - | - | - | - | - | - | 11 | 0 | 11 | 0 | PCR | *wsp* | (18) |
| China | 5 | 26 | 2014 | 693 | - | - | - | - | - | - | - | - | - | - | 647 | 16 | 70 | 561 | PCR | *wsp*, 16 S rDNA | (15) |
| HK, China | 5 | 26 | 2018–2019 | 1582 | - | - | - | - | - | - | - | - | - | - | 1310 | 358 | 375 | 577 | PCR | 12S rDNA | (75) |
| Cameroon | 4 | 23 | 2017–2021 | 155 | - | - | - | - | - | - | - | - | - | - | 8 | - | - | - | qPCR | 16S rRNA | (78) |
| China | 5 | 20 | 2013–2014 | 341 | 83 | 82 | - | - | - | 258 | 249 | - | - | - | 331 | - | - | - | qPCR | *wsp*, 16 S rRNA | (81) |
| Multiple | - |  | 2022–2023 | 104 | - | - | - | - | - | - | - | - | - | - | 104 | - | - | - | qPCR | 16 S rRNA | (86) |
| Greece | 1.30 | 27 | 2021 | 105 | 31 | 28 | 0 | 12 | 16 | 74 | 72 | 0 | 4 | 68 | 100 | 0 | 16 | 84 | Multiplex PCR | *wsp* | (87) |
| Sri Lanka | 825 | 27 | - | 127 | - | - | - | - | - | - | - | - | - | - | 62 | - | - | - | PCR | *wsp* | (72) |
| Odisha, India | 72 | 21 | 2014-2016 | 323 | 71 | 60 | - | - | - | 252 | 222 | - | - | - | 282 | - | - | - | PCR | *wsp* | (79) |
| Hainan, China | 5 | 20 | 2020-2021 | 90 | - | - | - | - | - | - | - | - | - | - | 78 | - | - | - | PCR | *wsp* | (85) |
| Yucatan, Mexico | 1786 | 18 | 2019 | 45 | 16 | 7 | - | - |  | 29 | 11 |  | - | - | 18 | - | - | - | PCR | 16 S rRNA | (76) |
| Orissa, India | 72 | 21 | 2010-2012 | 1291 | 689 | 681 | - | - | - | 602 | 600 | - | - | - | 1281 | - | - | - | PCR | *wsp* | (67) |
| Cemeteries, Southern Mexico | 1786 | 18 | 2015 | 108 | - | - | - | - | - | - | - | - | - | - | 41 | - | - | - | PCR | *wsp* | (77) |
| Singapore | 990 | 28 | 2018-2019 | 37 | - | - | - | - | - | - | - | - | - | - | 21 | - | - | - | PCR | *wsp* | (74) |
| Analamanga, Madagascar | 2 | 20 | 2008 | 685 | 272 | 272 | - | 6 | - | 413 | 413 | - | 4 | 409 | 685 | - | - | - | PCR | *wsp* | (66) |
| KL, Malaysia | 2358 | 26 | 2012-2013 | 286 | 74 | 74 | 0 | 5 | 65 | 212 | 212 | 3 | 1 | 197 | 286 | 3 | 6 | 262 | PCR | *wsp* | (69) |
| Pernambuco, Brazil | 18 | 26 | - | 143 | 58 | 48 | - | - | - | 85 | 83 | - | - | - | 131 | - | - | 96 | PCR | *ftsZ* and *wsp* | (65) |
| Guangzhou, China | 5 | 20 | - | 223 | 113 | 113 | 9 | 0 | 104 | 110 | 110 | 0 | 1 | 109 | 223 | 9 | 1 | 213 | PCR | *wsp* | (56) |
| Valencia, Spain | 3 | 27 | 2019 | 50 | 25 | 23 | 1 | 9 | 13 | 25 | 24 | 3 | 0 | 21 | 47 | 4 | 9 | 34 | PCR | *wsp*, 16S rRNA | (83) |
| India | 72 | 21 | 2011 | 57 | - | - | - | - | - | - | - | - | - | - | 57 | 0 | 0 | 57 | PCR | *wsp* | (68) |
| Eldorado, Argentina | 5040 | 18 | - | 408 | - | - | - | - | - | - | - | - | - | - | 312 | 1 | 97 | 214 | PCR | *wsp* | (82) |

Two-thirds of the studies (24 of 36) had a low to moderate risk of bias, while close to one-third (12 of 36) had a high risk of bias according to the JBI critical appraisal tool (63)

### Natural occurrence of *Wolbachia* in *Aedes* mosquitoes

Random-effects model estimation was used in the meta-analysis, and the overall occurrence of *Wolbachia* in field-collected *Aedes* mosquitoes worldwide was 57.7% (95% CI: 41.0–72.8%), ranging from 6.0% (95% CI: 2.6–13.1%) in *Ae. aegypti* to 87.1% (95% CI: 78.0–92.8%) in *Ae. albopictus*. There was no evidence of publication bias (Kendall’s tau *p* = 0.85601), and between-study heterogeneity was significantly high (*I*^*2*^ = 98.81; Q test p < 0.001). The high degree of heterogeneity was related to the mosquito species and dengue incidence (*p* <0.03) (Table **2**).

**Table 2.** Meta-regression of covariates with random-effect model.

| Covariate | Category | Coefficient | SE | 95% CI for Logit event rate |  | p-value |
| --- | --- | --- | --- | --- | --- | --- |
|  |  |  |  | Lower | Upper |  |
| Intercept |  | 2.8304 | 0.4820 | 1.8858 | 3.7750 | 0.0000 |
| Species | *Ae. albopictus* | Reference |  |  |  |  |
|  | *Ae. aegypti* | -5.5134 | 0.6469 | -6.7814 | -4.2454 | 0.0000 |
| Dengue incidence<sup>s</sup> |  | -0.0006 | 0.0003 | -0.0011 | -0.0001 | 0.0281 |
| Area temperature* | Normal | Reference |  |  |  |  |
|  | High | -0.9877 | 0.6051 | -2.736 | 0.1982 | 0.1026 |

<sup>s</sup>The annual incidence of dengue per million individuals; \* Temperature: High (> 25 °C) and Normal (≤ 25 °C)

Across the continent, Asia had the highest percentage of *Wolbachia* infection in *Ae. aegypti* (7.1%), followed by Europe (5.0%), North America (1.9%), and Africa (0.7%) (Fig. **2**). Similarly, Asia had the highest prevalence of *Wolbachia* in *Ae. albopictus* (95.5%), followed by Europe (94.8%), North America (91.6%), South America (85.2%), and Africa (71.6%) (Fig. **3**).

**Fig. 2.**
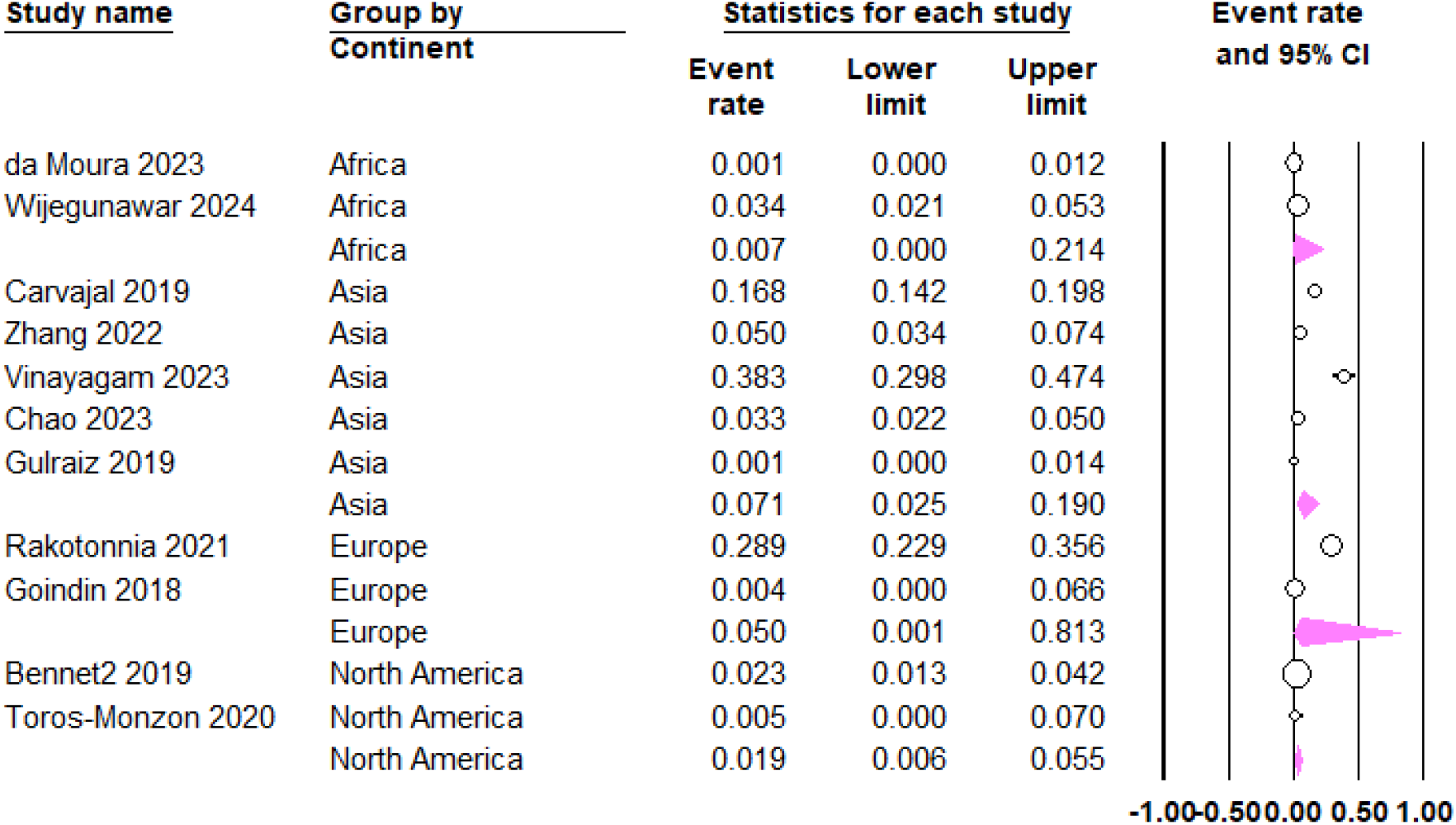
Prevalence of natural *Wolbachia* infection in field-collected *Ae. aegypti* by continent.

**Fig. 3.**
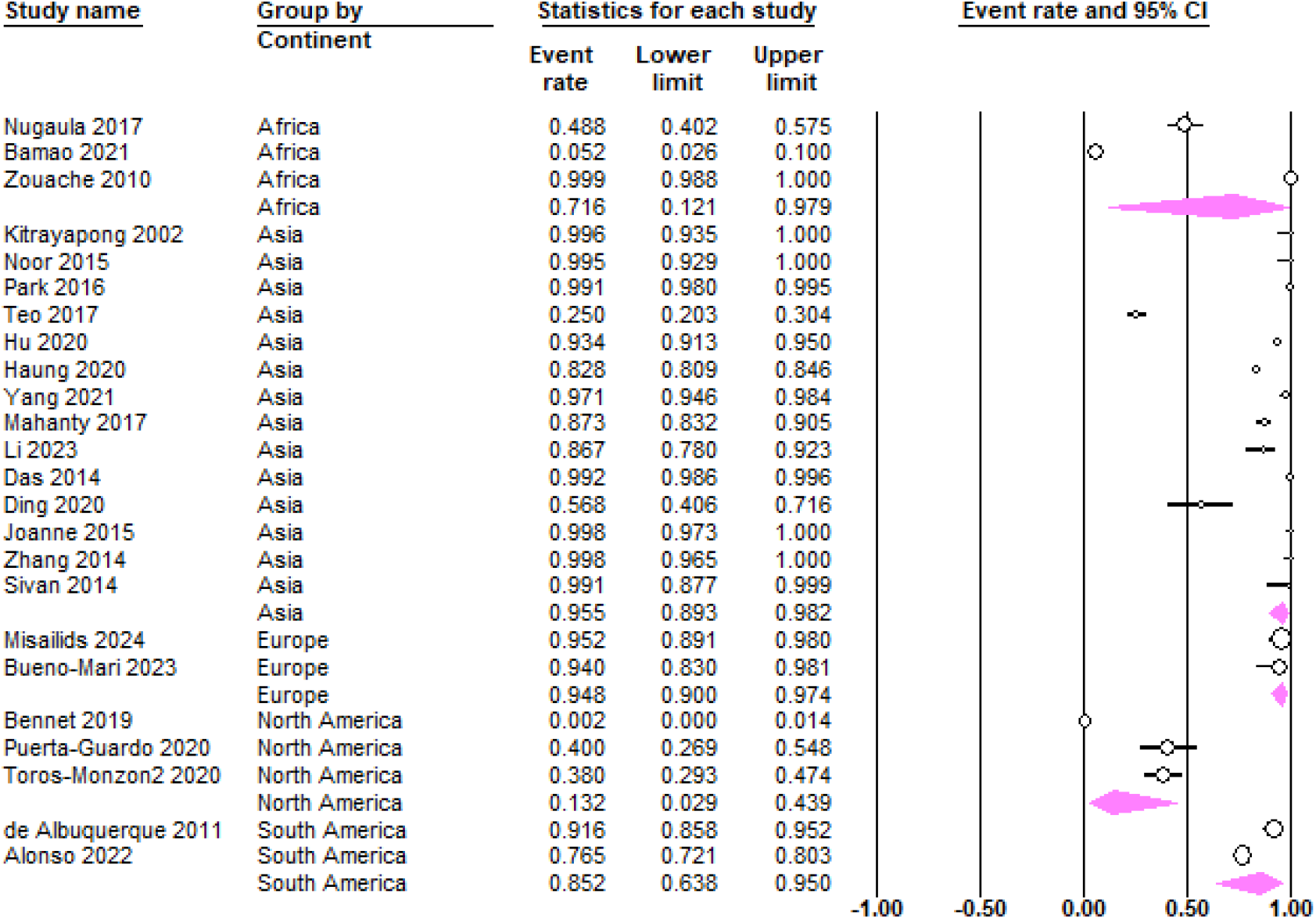
Prevalence of natural *Wolbachia* infection in field-collected *Ae. albopictus* by continent.

The highest prevalence of *Wolbachia* in *Ae. aegypti* was seen in India (38.3%) (20) followed by France (28.9%) (80) and the Philippines (26) (Fig. **2**). However, no evidence of *Wolbachia* infections was reported in *Ae. aegypti* in a multi-continent study (59), in Pakistan (89), Cabo Verde (84), France (61), and Mexico (77) (Table 1). On the contrary, the majority (over 70%) of field-collected *Ae. albopictus* harbours the endosymbiotic bacteria in Thailand (64), Madagascar (66), Malaysia (69, 70), Korea (71), China (15, 56, 75, 81, 85), India (67, 68, 79), Spain (83), and Greece (87), while less than half of *Ae. albopictus* mosquitoes were naturally infected with *Wolbachia* in Cameroon (78), Sri Lanka (72), Malaysia (21), Panama (18), and Mexico (76, 77) (76, 77).

### Factors associated with *Wolbachia* prevalence in *Aedes* mosquitoes

The prevalence of *Wolbachia* infection in *Ae. aegypti* was significantly higher among females than males (OR, 1.72; 95% CI, 1.01, 2.92, *p* = 0.046) (Fig. **4**), while there was no difference between males and females in *Ae. albopictus* (OR, 0.54; 95% CI, 0.26, 1.12, *p* = 0.098) (Fig. **5**).

The prevalence rate of *Wolbachia* was more than six times lower in *Ae. aegypti* than in *Ae. albopictus* (*p* <0.0001). Besides, the overall *Wolbachia* prevalence among *Aedes* mosquitoes was associated with dengue incidence (p <0.03) (Table **2**).

**Fig. 4.**
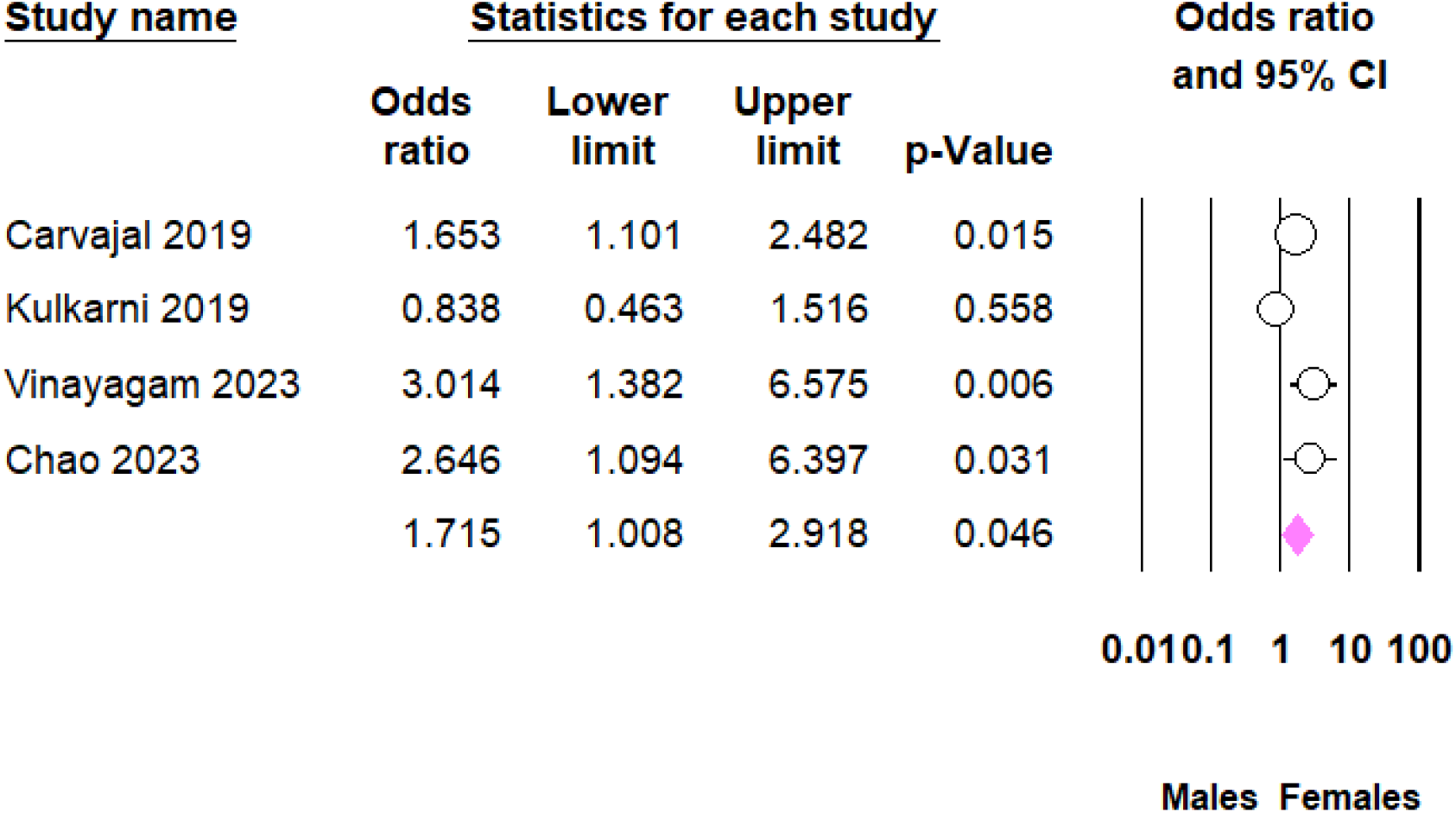
Host sex difference in the prevalence of *Wolbachia* infection in *Ae. aegypti*.

**Fig. 5.**
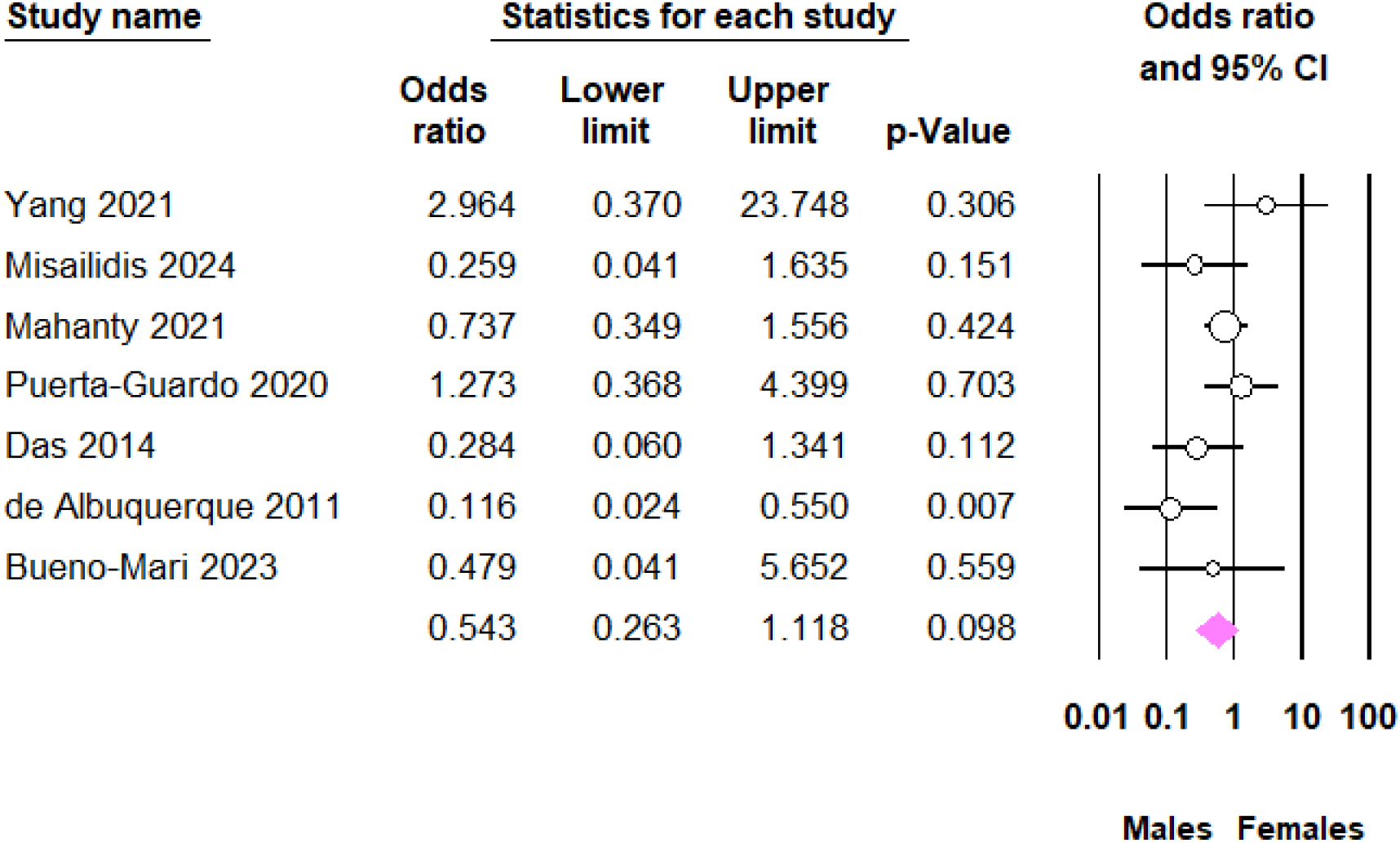
Host sex difference in the prevalence of *Wolbachia* infection in *Ae. albopictus*.

More specifically, *Wolbachia* prevalence was negatively associated with dengue incidence in *Ae. albopictus (p* <0.01) (Table **3**), but this was not related to dengue incidence in *Ae. aegypti* (*p* >0.05) (Table **4**). On the contrary, *Wolbachia* prevalence was negatively associated with area temperature in *Ae. aegypti (p* <0.001) (Table **4**), but this was not related to area temperature in *Ae. albopictus* (*p* >0.05) (Table **3**).

**Table 3.**
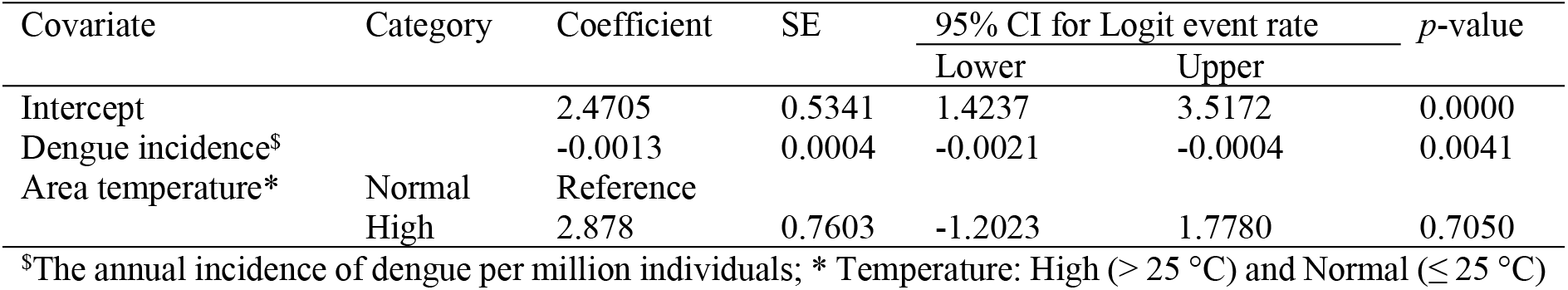
Meta-regression of covariates of *Wolbachia* infection in *Ae. albopictus* with random-effect model.

**Table 4.** Meta-regression of covariates of *Wolbachia* infection in *Ae. aegypti* with random-effect model.

| Covariate | Category | Coefficient | SE | 95% CI for Logit event rate |  | p-value |
| --- | --- | --- | --- | --- | --- | --- |
|  |  |  |  | Lower | Upper |  |
| Intercept |  | -2.6153 | 0.5761 | -3.7445 | -1.4861 | 0.0000 |
| Dengue incidence<sup>s</sup> |  | 0.0009 | 0.0005 | -0.0002 | 0.0019 | 0.0984 |
| Area temperature* | Normal | Reference |  |  |  |  |
|  | High | -2.5736 | -3.9410 | -1.2061 | -3.69 | 0.0002 |

<sup>s</sup>The annual incidence of dengue per million individuals; \* Temperature: High ( $> 25^{\circ}\text{C}$ ) and Normal ( $\leq 25^{\circ}\text{C}$ )

## Discussion

Pathogen-blocking and reproductive changes in arthropods, such as parthenogenesis, feminization, male death, cytoplasmic incompatibility, and female infertility, can be caused by *Wolbachia* (14), an endosymbiotic gram-negative bacterium recently introduced as an environmentally friendly novel dengue biocontrol agent (6). Therefore, it is vital to determine the natural occurrence of *Wolbachia* in *Aedes* mosquitoes. To the authors’ knowledge, this is the first meta-analysis to document the natural occurrence of *Wolbachia* in *Ae. aegypti* and *Ae. albopictus*. About 16,997 field-collected *Aedes* mosquitoes (7,531 *Ae. aegypti* and 9,466 *Ae. albopictus*) were included in 36 studies conducted in different countries.

Based on the meta-analysis, the pooled prevalence of *Wolbachia* in *Ades* mosquitoes worldwide was 57.7% (95% CI: 41.0–72.8%), ranging from 6.0% (95% CI: 2.6–13.1%) in *Ae. aegypti* to 87.1% (95% CI: 78.0–92.8%) in *Ae. albopictus*.

Among the continents, Asia had the highest percentage of *Wolbachia* infection in *Ae. aegypti* (7.1%), followed by Europe (5.0%). Similarly, Asia had the highest prevalence of *Wolbachia* in *Ae. albopictus* (95.5%), followed by Europe (94.8%), North America (91.6%), South America (85.2%), and Africa (71.6%). The variation could be attributed to differences in detection methods (28); environmental factors such as temperature (55); and the genetic variation in the mosquito populations (91).

Factors associated with the overall *Wolbachia* prevalence include mosquito species (*p* <0.0001) and dengue incidence (*p* <0.03), i.e., the natural occurrence of *Wolbachia* in *Ae. albopictus* was six times higher than in *Ae. aegypti*, and dengue incidence was negatively related to *Wolbachia* prevalence. While the natural infection of *Ae. albopictus* is more common, *Wolbachia* in *Ae. aegypti* is often found in low density and high sequence diversity, and its presence could reflect contamination rather than a true signal (17, 92).

Species wise, infection rates in *Ae. aegypti* were significantly higher among females than males (OR = 1.72; 95% CI = 1.01, 2.92, *p* = 0.046), while there was no difference between males and females in *Ae. albopictus* (*p* = 0.098). Besides, *Wolbachia* infection rates in *Ae. albopictus* were inversely correlated with dengue incidence (β = −0.0013, *p* <0.01). This could be due to the viral blocking effect of the endosymbiont (93).

Furthermore, higher temperature was negatively associated with *Wolbachia* prevalence in *Ae. aegypti* (β = −2.5736, p <0.001). This could be due to the heat stress during larval development directly impacting the replication and transmission of the *Wolbachia* bacteria, leading to a lower bacterial load within the mosquito (94).

This study has several strengths and limitations. The study’s strengths were the large sample size and the lack of publication bias among the included studies; its weaknesses included limiting the study to only those published in English and finding evidence of significant heterogeneity among the studies.

## Conclusion

*Aedes* mosquitoes had a high prevalence of naturally occurring *Wolbachia*, but at varying rates between nations and continents, which was negatively correlated with dengue incidence. The identification of natural *Wolbachia* in *Ae. aegypti* may indicate contamination, even though *Ae. albopictus* infections are more common. Therefore, high-quality multi-centre studies on *Wolbachia* prevalence in *Aedes* mosquitoes are required to verify the above findings.

## Declarations

### Ethics approval, and consent to participate

Our study did not require ethical approval, as the data used have been published previously and are hence already in the public domain. Consent was not required when conducting a systematic review.

## Consent for publication

Not applicable.

## Availability of data and materials

The study data are available within the manuscript.

## Author contributions

TTG, under the supervision of PL, designed the study, wrote the statistical analysis plan, monitored the review process, interpreted the data, cleaned and analysed the data, and wrote the draft manuscript. TTG and VR assessed studies for inclusion. All authors have approved the final version.

## Acknowledgements

The corresponding author would like to thank Prof. Ary Hoffmann and Dr. Perran Stott-Ross from the University of Melbourne for guidance during the review process.

